# Fitness cost of vancomycin-resistant *Enterococcus faecium* plasmids associated with hospital infection outbreaks

**DOI:** 10.1101/2021.01.27.428397

**Authors:** Ana P. Tedim, Val F. Lanza, Concepción M. Rodríguez, Ana R. Freitas, Carla Novais, Luísa Peixe, Fernando Baquero, Teresa M. Coque

**Affiliations:** Department of Microbiology, University Hospital Ramón y Cajal, Madrid, Spain; UCIBIO/REQUIMTE. Department of Biological Sciences, Microbiology Laboratory, Faculty of Pharmacy, University of Porto, Porto, Portugal; Centres for Biomedical Research in the Epidemiology and Public Health Network (CIBER-ESP)

**Keywords:** Vancomycin resistant Enterococcus, Fitness cost, Tn*1546*, vancomycin resistance plasmids, Enterococcus faecium

## Abstract

**Background:** Vancomycin resistance is mostly associated with *Enterococcus faecium* due to Tn*1546*-*vanA* located on narrow- and broad-host plasmids of various families. The study’s aim was to analyse the effects of acquiring Tn*1546*-plasmids with proven epidemicity in different bacterial host backgrounds.

**Methods:** Widespread Tn*1546*-plasmids of different families RepA_N (n=5), Inc18 (n=4) and/or pHTβ (n=1), and prototype plasmids RepA_N (pRUM) and Inc18 (pRE25, pIP501) were analysed. Plasmid transferability and fitness cost were assessed using *E. faecium* (GE1, 64/3) and *Enterococcus faecalis* (JH2-2/FA202/UV202) recipient strains. Growth curves (Bioscreen C) and Relative Growth Rates were obtained in presence/absence of vancomycin. Plasmid stability was analysed (300 generations). Whole genome sequencing (Illumina-MiSeq) of non-evolved and evolved strains (GE1/64/3 transconjugants, n=49) was performed. SNP calling (breseq software) of non-evolved strains was used for comparison.

**Results:** All plasmids were successfully transferred to different *E. faecium* clonal backgrounds. Most Tn*1546*-plasmids and Inc18 and RepA_N prototypes reduced host fitness (−2%-18%) while the cost of Tn*1546* expression varied according to the Tn*1546*-variant and the recipient strain (9-49%). Stability of Tn*1546*-plasmids was documented in all cases, often with loss of phenotypic resistance and/or partial plasmid deletions. Point mutations and/or indels associated with essential bacterial functions were observed on the chromosome of evolved strains, some of them linked to increased fitness.

**Conclusions:** The stability of *E. faecium* Tn*1546*-plasmids in the absence of selective pressure and the high intra-species conjugation rates might explain the persistence of vancomycin resistance in *E. faecium* populations despite the significant burden they might impose on bacterial host strains.

## INTRODUCTION

Vancomycin Resistant Enterococci (VRE) have increasingly been reported worldwide since their first description in the late 1980s,^1^ and are mostly associated with *Enterococcus faecium*. The most frequent vancomycin-resistant genotypes are *van*A (Tn*1546*) and *van*B (Tn*1549/5382*), which have followed various distinct dissemination pathways through plasmids and chromosomal conjugative elements, respectively.^2^

Tn*1546* is mostly located on transferable plasmids,^1,3^ which have greatly contributed to the successful spread of glycopeptide resistance in hospitals and farms. In fact, Tn*1546* is overrepresented in *E. faecium* lineages^1^ predominant in hospitalised patients (clade A1) and animals (clade A2). Strains from healthy humans fall into a separate clade B which remains mostly susceptible to glycopeptides and other antibiotics.^1,4^ Various studies have demonstrated the key contribution of plasmids in the evolution and adaptation of *E. faecium* to different challenges including antibiotics.^5^ Tn*1546*-carrying plasmids contain one or more genes coding for replication initiator proteins (*rep*) of the Inc18 (e.g. pIP501 and pRE25), RepA_N (e.g. pRUM and pLG1) and to a lesser extent, pHTβ plasmid families.^1,6,7^ In a previous study, our group suggested that the emergence of particular RepA_N and Inc18 Tn*1546*-carrying plasmids in the US and Europe in the late 1990s (pRUM and pIP186, respectively) could have contributed to the further spread of Tn*1546* in American hospitals and European farms.^1^

A few studies have shown that Tn*1546*-carrying plasmids reduce the fitness of their bacterial host. However, such plasmids were apparently stable in these hosts as the initial fitness cost was rapidly decreased by compensatory changes in sequential evolution experiments.^8,9^ Unfortunately, these studies were limited to a single recipient strain and/or poorly characterised plasmids.^8–13^

The aim of this study was to determine the fitness cost imposed by the acquisition of field enterococcal plasmids associated with documented hospital infection outbreaks worldwide.^1^ We also studied the intrinsic fitness of *E. faecium* strains of various clades to further understand the maintenance or propensity of Tn*1546* to persist in specific populations. These *E. faecium* populations are probably those with a higher intrinsic fitness, and/or where the uptake of plasmids harbouring particular Tn*1546* variants might impose a lower fitness cost, in the absence of or during antibiotic exposure.

## MATERIAL AND METHODS

### Bacterial strains and plasmids

We analysed seven *E. faecium* strains harbouring transferable Tn*1546-*carrying emblematic plasmids known for their apparent epidemicity and persistence in hospital settings were analysed.^1,14^ These plasmids present *rep* genes associated with various plasmid families, Inc18 (4 rep_1/pIP501_ and 3 rep_2/pRE25_), RepA_N (3 rep_17/pRUM_ and 2 rep_20/pLG1_), and pHTβ (1 rep_22/pHTβ_). For comparison, we also studied prototype plasmids for Inc18 [pIP501-rep_1/pIP501_ and pRE25-rep_1/pIP501_+rep_2/pRE25_ (GenBank NC_008445)] and RepA_N [pRUM (GenBank NC_005000)] plasmid families, from which current enterococcal plasmid chimeras seem to derive.^15–17^ We used rifampicin and fusidic acid-resistant *E. faecium* GE1 (ST515) and 64/3 (ST21), and *E. faecalis* JH2-2, FA202 and UV202 (ST8) as recipient strains. The characteristics of the strains and plasmid analysed are shown in Figure 1 and Table S1.

**Figure 1.**
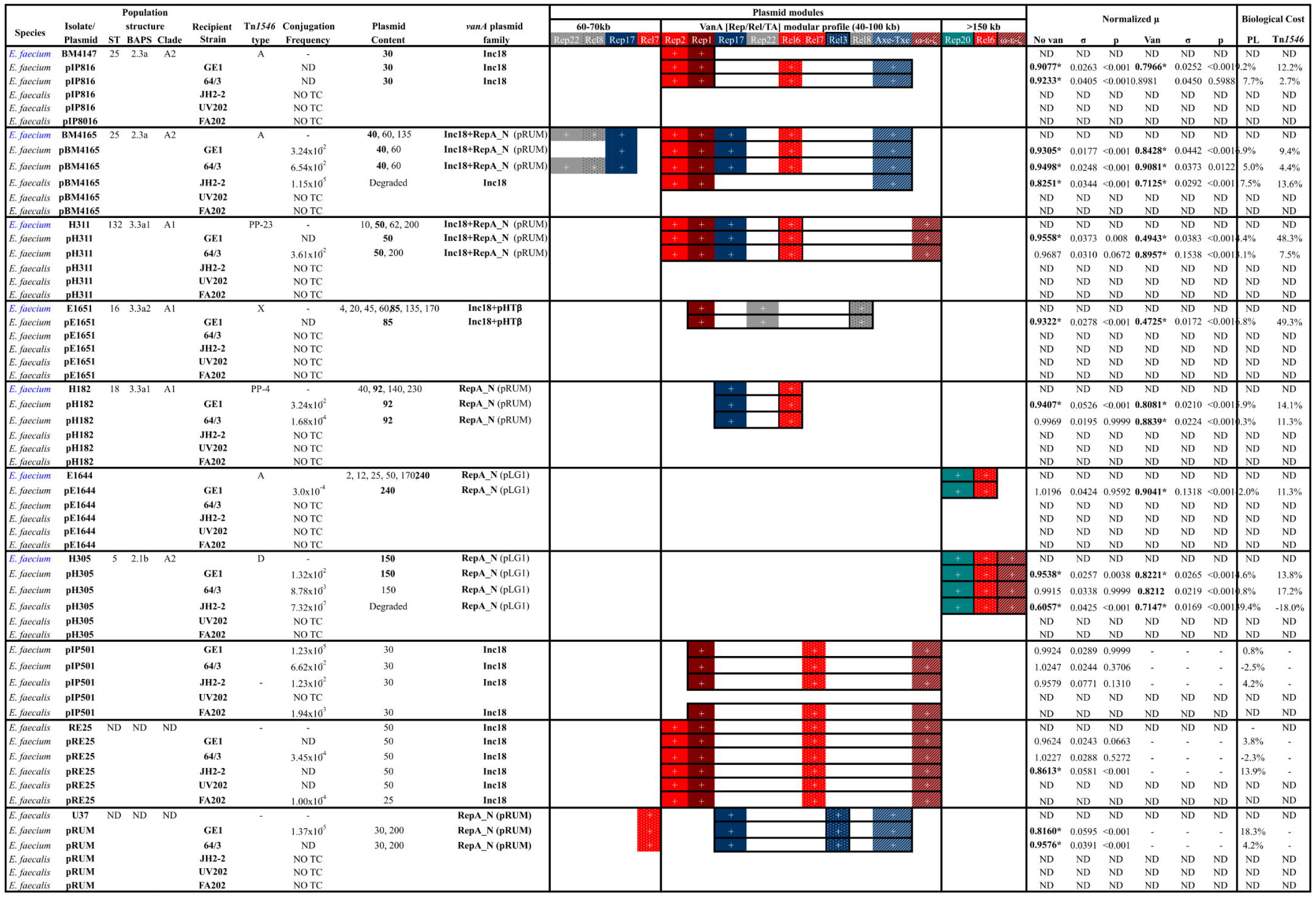
The Tn*1546-vanA* and prototype plasmids employed in this study. Rep1, *rep*_1/pIP501_; Rep2, *rep*_2/pRE25_; Rep17, *rep*_17/pRUM_; Rep22, *rep*_22/pHTβ_; Rel3, *rel*_3/pRUM_; Rel6, *rel*_6/pEF1_; Rel7, *rel*_7/pIP501_; Rel8, *rel*_8/pHTβ_; ωεζ, Toxin-Antitoxin system of plasmid pSM19035; Axe-Txe, Toxin-Antitoxin system of plasmid pRUM. *statistically significant. Plasmid in bold - plasmid carrying Tn*1546-vanA*. Abbreviations: ND, Not Determined; NO TC, No transconjugant was obtained; No Van, No vancomycin induction; Van, Vancomycin induction; PL, Plasmid.

### Plasmid analysis and transferability

We determined the plasmid content by S*1*-nuclease pulsed field gel electrophoresis (PFGE) and characterisation of *rep* genes, relaxases (*rel*) and maintenance systems (partition and toxin-antitoxin systems) by PCR, hybridisation and further sequencing.^1,18,19^

We determined plasmid transferability by filter mating experiments at a 1:1 donor-recipient ratio and overnight incubation, as previously described.^20^ Transconjugants were selected on Brain Heart Infusion (BHI) agar plates containing fusidic acid (20μg/mL), rifampicin (30μg/mL) and vancomycin (6μg/mL) or erythromycin (20μg/mL) after a 24h incubation at 37°C.^20,21^ We calculated the conjugation frequencies as the proportion of transconjugants colony-forming units (CFUs) per recipient CFUs and confirmed the transconjugants by PCR detection of *vanA* and *erm(B)* genes,^22,23^ and by *Sma*I-PFGE (Takara Bio Inc., Shiga, Japan).^24^

### Growth kinetics

We analysed the growth kinetics of 63 well characterised field strains of *E. faecium* (Supplementary Table S1), recipient *E. faecium* (GE1 and 64/3) and *E. faecalis* (JH2-2) and transconjugants harbouring Tn*1546*-carrying plasmids using the Bioscreen C plate reader (ThermoLab Systems, Vantaa, Finland) adapting the method described by Foucault *et al.*^10^ Briefly, one colony per strain was grown overnight (18h) at 37°C in BHI broth with and without induction (6mg/L of vancomycin). Overnight cultures were diluted 1:1000 into fresh BHI broth, approximately 10^5^ CFUs/mL, and 300μl of this bacterial suspension was transferred into a 100-well microplate. We measure the optical density (OD) at 600 nm every 15 min for 20 h, Keeping the temperature at 37°C. To ensure culture optical homogeneity, the plates were shacked for 10s before each OD measurement. For each strain, 5 biological replicates were assayed in duplicate in each experiment (10 readings *per* strain *per* experiment). We performed two independent experiments were performed for all strains analysed,^10^ maintaining the induction or non-induction conditions throughout the entire experiment. We employed variants of this procedure to analyse the possible changes in the *E. faecium* growth dynamics at 42°C, given this is the body temperature of poultry, a known reservoir of VRE.

We determined the growth rates (μ) in the interval estimated to be the exponential phase using the GrowthRates 2.1 program.^25^ To better compare the results between experiments, we performed the normalisation using the mean growth rate of the plasmid free recipient strains (GE1, 64/3 or JH2-2) per experiment. We determined the fitness cost of the plasmids and the fitness cost of Tn*1546* expression by calculating the Relative Growth Rate influenced by plasmid (RGR_PL_) and Relative Growth Rate influenced by transposon expression (RGR_Tn_) using the following formulas:

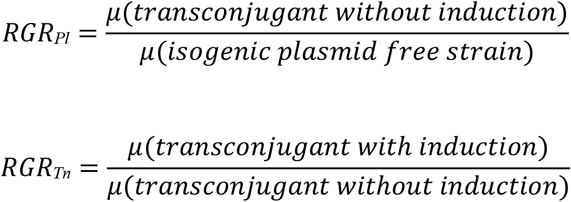

### Plasmid stability

We inoculated three independent colonies *per* strain (recipients and transconjugants, see Supplementary Figure S1) into 5 mL of BHI broth (1 tube *per* colony *per* strain) and incubated them at 37°C overnight (22-24 h). We subsequently, diluted the cultures 1:1000 into fresh BHI broth and incubated them at 37°C overnight (22-24 h). We plated the bacterial cultures at 0, 100, 200 and 300 generations onto BHI agar plates and then randomly picked 100 colonies of each plate onto BHI agar plates supplemented or not with the appropriate antibiotic (6 μg/mL of vancomycin for strains containing pH182, pH311 and pBM4165; 20 μg/mL of erythromycin for strains containing pIP501, pRE25 and pRUM).^15–17^ We initially calculated the plasmid loss frequency by determining the ratio of susceptible colonies to the total number of colonies and further confirmed antibiotic susceptibility (stability/loss) at phenotypic (disk-diffusion) and genotypic (PCR of *vanA* and *ermB* genes) levels as previously described.^22,23^ We further checked the presence/absence of plasmid by S*1*-PFGE and by confirming the presence of all plasmid modules previously observed in each of the plasmids studied, as described above (Table 1).

### Genome analysis of evolved strains

We performed whole genome sequencing of 49 transconjugants, 24 *E. faecium* GE1 (7 non-evolved and 17 evolved strains) and 25 *E. faecium* 64/3 (7 non-evolved and 18 evolved strains) using an Ilumina MiSeq platform to obtain 100-200 bp paired-end reads (Supplementary Figure S1). The DNA extraction was performed using the Promega Wizard Genomic DNA Purification kit (Promega, WI, USA) according to the manufacturer’s instructions. For the non-evolved strains, we revised the reads and corrected the sequencing errors with Lighter software,^26^ estimating the best k-mer length with KmerGenie software,^27^ and performing the final assembly with SPAdes.^28^1 We performed an single nucleotide polymorphism analysis of the evolved strains against the non-evolved strains using the Breseq v0.26.1 pipeline (http://barricklab.org/twiki/bin/view/Lab/ToolsBacterialGenomeResequencing).^29^. We classified the protein functions according to the database of Clusters of Orthologous Groups of proteins (COGs) and performed a plasmid analysis of the non-evolved strains using the plasmid reconstruction tool PLACNET (Figures S2 and S3) to determine the plasmid content in these strains and identify the location of mutations in the evolved strains.^30^

We also performed growth kinetics experiments for these evolved strains as described above. The non-evolved strains for each background and plasmid, as well as the plasmid-free strains (GE1 and 64/3) were included to normalise the results. We calculated the relative growth rates and costs using the following equations:

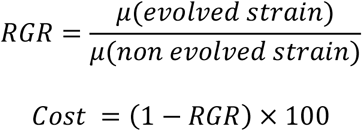

### Statistical analysis

We calculated the statistics using the R analytical Studio package,^31^ employing analysis of variance and Tukey’s honest significance tests (for multiple *all in all* comparison). We employed the chi-squared test when comparing two populations. Values of *p*<0.01 were considered statistically significant.

## RESULTS

### Fitness cost of the plasmids carrying Tn*1546* (*vanA*) in *E. faecium* and *E. faecalis*

Tn*1546-van*A-carrying plasmids and prototype plasmids of the Inc18, RepA_N plasmid (pRUM/pLG1) and pHTβ families imposed different fitness costs on distinct clonal backgrounds (Figures 1 and 2). With the exception of pE1644 (rep_20/pLG1_+rel_6_, fitness gain of 2.0%, p=0.9658), all tested plasmids imposed a fitness cost on *E. faecium* GE1 (n=7, ranging from 4.4% to 9.2%) and, to a lesser extent, in *E. faecium* 64/3 (n=5, 0.3-7.7%). The two *E. faecium* plasmids that transferred to *E. faecalis* JH2-2 (pBM4165 and pH305) imposed a high fitness cost on this strain (17.5% and 39.4%, respectively).

**Figure 2.**
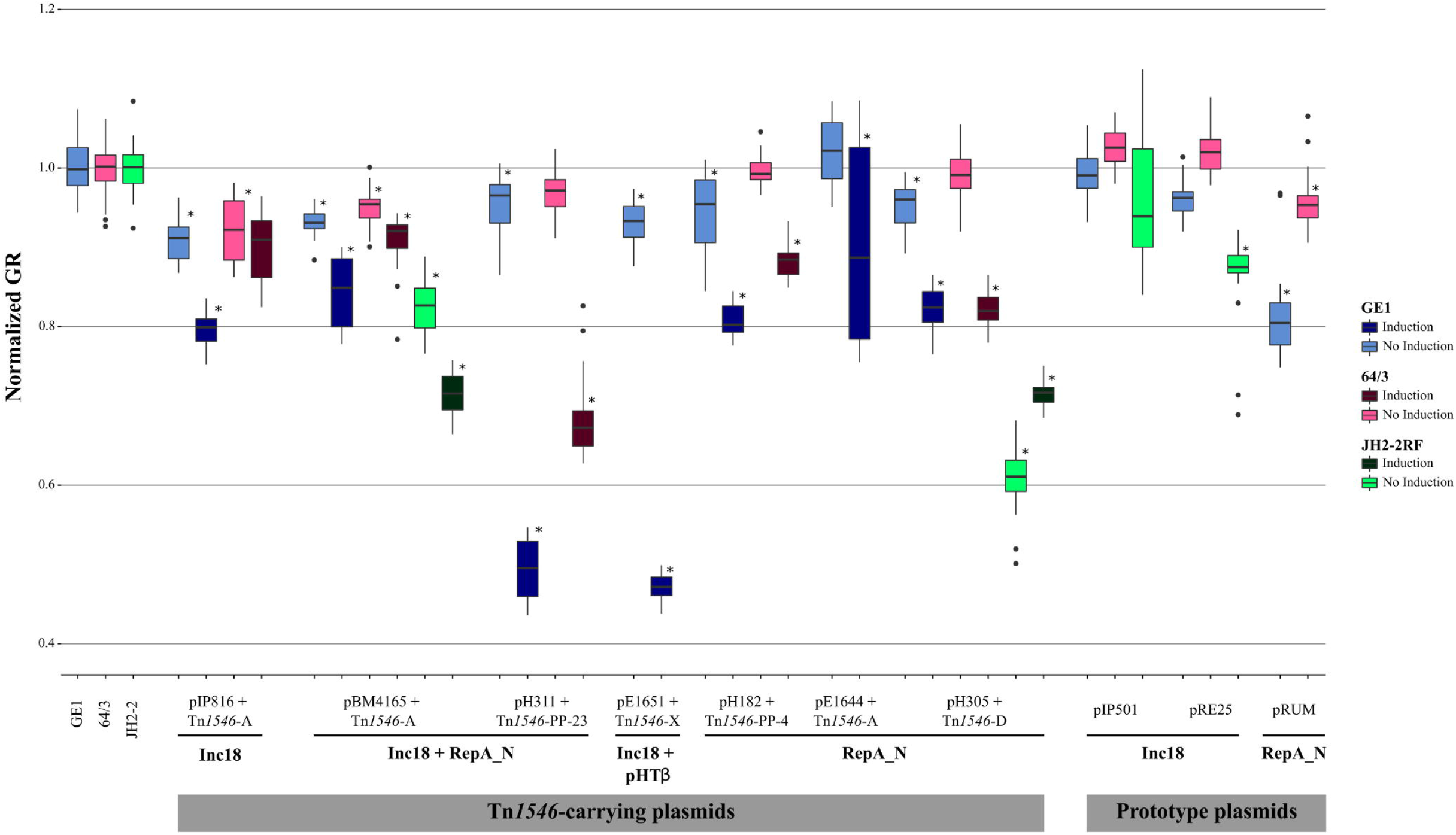
Box and whiskers plot representing the normalised growth rates of *E. faecium* Tn*1546*-*vanA* carrying plasmids with and without vancomycin induction and prototype plasmids (pRE25, pIP501 and pRUM). Plasmid and Tn*1546*-*vanA* expression were tested in different backgrounds (*E. faecium* GE1 and 64/3 and *E. faecalis* JH2-2) when plasmids conjugated into those backgrounds. Plasmids are classified according to the replication initiation protein plasmid family and Tn*1546* type. *statistically significant. Abbreviations: GR, growth rate.

#### Fitness of Inc18 family plasmids

pIP816 (rep_1/pIP501_+rep_2/pRE25_+rel_6_+TA_Axe-Txe_), the first *vanA* plasmid described in 1986,^1,32^ exhibited the highest fitness cost values in both *E. faecium* GE1 (9.2%) and 64/3 (7.7%). Other Inc18 and Inc18-mosaic plasmids isolated afterwards (pE1651, rep_1/pIP501_+rep_22/pHTβ_+rel_8/pHTβ_; pBM4165, rep_1/pIP501_+rep_2/pRE25_+rep_17/pRUM_+rel_6_+TA_Axe-Txe_; and pH311, rep_1/pIP501_+rep_2/pRE25_+rep_17/pRUM_+rel_6_+TA_εζ_) also reduced the fitness of *E. faecium* GE1 (6.8%, 6.9% and 4.4%, respectively, but the reduction was not statistically significant). Conversely, the Inc18 prototypes pIP501 and pRE25 did not impose a significant fitness cost on the *E. faecium* strains studied, although the carriage of pRE25 provoked a growth reduction on *E. faecalis* JH2-2 (13.9%). Aside from pIP816, the only plasmid that imposed a significant fitness cost on *E. faecium* 64/3 was a mosaic Inc18+RepA_N plasmid (pBM4165, 5.0%), this plasmid as reported in Europe in the late 1980s (Figures 1, 2 and Table S1).

#### Fitness of RepA_N family plasmids

The RepA_N-pRUM plasmids studied (pH182, rep_17/pRUM_+rel_6_) also imposed a significant fitness cost on *E. faecium* GE1 (5.9%), as did the pRUM prototype in both *E. faecium* backgrounds (GE1, 18.4%; 64/3, 4.2%). RepA_N-pLG1 plasmid (pH305, rep_22/pLG1_+rel_6_+TA_εζ_) reduced the fitness of *E. faecium* GE1 by 4.6% but did not significantly reduce that of *E. faecium* 64/3.

### Fitness cost of Tn*1546* expression in isogenic *E. faecium* and *E. faecalis* backgrounds

The fitness cost of Tn*1546* expression varied according with the transposon variant, and the clonal and plasmid backgrounds in which they were located (Figures 1 and 2).

The expression of the Tn*1546* variant A of mosaic plasmid pBM4165 (Inc18+RepA_N) imposed a fitness cost of 9.4% on *E. faecium* GE1, 13.6% in *E. faecalis* JH2-2 and no fitness cost on *E. faecium* 64/3. The Tn*1546* variants PP-4 (containing an IS*Ef1* insertion in the *vanX-vanY* intergenic region), in pH182 (RepA_N), and variant D in plasmid pH305 (Inc18+RepA_N), imposed a similar cost on *E. faecium* GE1 and *E. faecium* 64/3 (14.1% *vs* 11.3% and 13.8% *vs* 17.2%, respectively). Interestingly, in the case of *E. faecalis* JH2-2, the expression of Tn*1546* variant D improves the growth rate of the background by 18.0% (Figures 1 and 2).

Other transposon variants exhibiting indels and duplications such as Tn*1546* variant PP-23 in pH311 (Inc18+RepA_N) and Tn*1546* variant X in pE1651 (Inc18+pHTβ) imposed a high fitness cost on GE1 (48.3% *vs* 49.3%) and 64/3 strains (29.1%, only analysed for pH311) (Figures 1 and 2).

### Plasmid transferability

The conjugation frequency varied among the plasmids and, for the same plasmid, between the recipient strains used. Most of *E. faecium* plasmids transferred at high frequencies (10^−2^ to 10^−4^) into all *E. faecium* recipients used but not to *E. faecalis*. Only two plasmids (pH305 and pBM4165) transferred to both *E. faecalis* and *E. faecium* (10^−7^ and 10^−5^, respectively) (Figure 1). While the Inc18 family prototype plasmids pRE25 and pIP501 were transferred from *E. faecalis* to most of the *E. faecalis* and *E. faecium* recipients used (10^−2^ to 10^−5^); the original pRUM plasmid from *E. faecium* was only successfully transferred to *E. faecium* recipients (approximately 10^−5^) (Figure 1).

### In-host long-term plasmid stability

*E. faecium* Tn*1546*-carrying plasmids were highly stable up to 300 generations in both *E. faecium* GE1 and *E. faecium* 64/3, although the presence of these plasmids was not always associated with a resistance phenotype. In general, all plasmids were more stable in *E. faecium* 64/3 than in GE1 (Figure 3).

**Figure 3.**
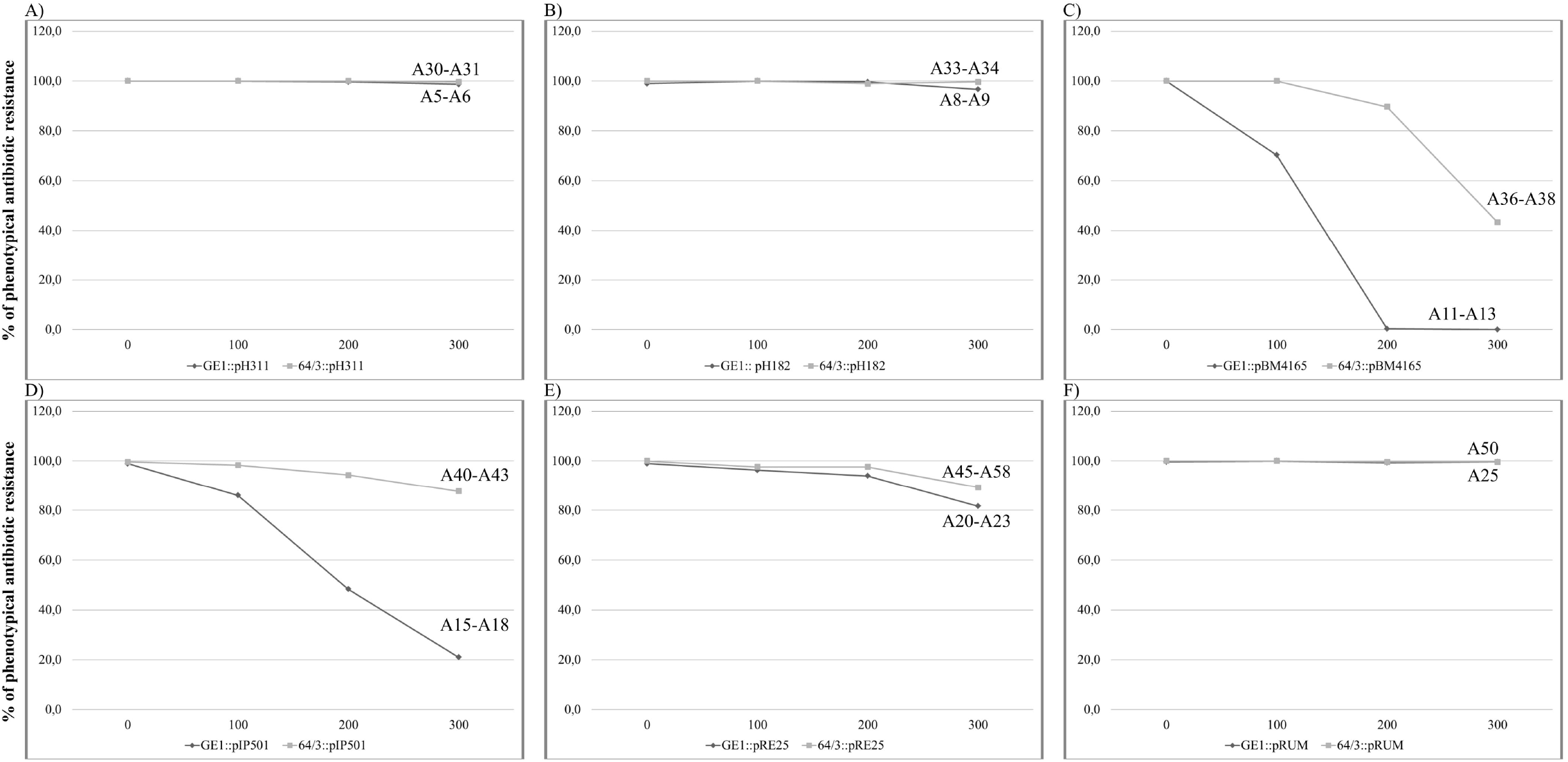
Percentage of antibiotic resistant phenotype loss of Tn*1546-vanA* carrying plasmids in *E. faecium* GE1RF and 64/3. Antibiotic resistance does not always correspond to plasmid loss. A) Antibiotic resistance phenotype loss for pH311 (A5 - GE1::pH311_300_::*vanA*; A6 - GE1::pH311_300_::Δ*vanA*; A30 - 64/3::pH311_300_::*vanA*; A31 - 64/3::pH311_300_::Δ*vanA*); B) Antibiotic resistance phenotype loss for pH182 (A8 - GE1::pH182_300_::*vanA*; A9 - GE1::pH182_300_::Δ*vanA*; A33 - 64/3::pH182_300_::*vanA*; A34 - 64/3::pH182_300_::Δ*vanA*); C) Antibiotic resistance phenotype loss for pBM4165 (A11 - GE1::pBM4165_100_::*vanA*; A12 - GE1::pBM4165_100_::Δ*vanA*; A13 - GE1::pBM4165_300_::Δ*vanA*; A36 - 64/3::pBM4165_100_::*vanA*; A37 - 64/3::pBM4165_300_::*vanA*; A38 - 64/3::pBM4165_300_::Δ*vanA*); D) Antibiotic resistance phenotype loss for pIP501 (A15 - GE1::pIP501_100_::*ermB*; A16 - GE1::pIP501_100_::Δ*ermB*; A17 - GE1::pIP501_300_::*ermB*; A18 - GE1::pIP501_300_::Δ*ermB*; A40 - 64/3::pIP501_100_::*ermB*; A41 - 64/3::pIP501_100_::Δ*ermB*; A42 - 64/3::pIP501_300_::*ermB*; A43 - 64/3::pIP501_300_::Δ*ermB*); E) Antibiotic resistance phenotype loss for pRE25 (A20 - GE1::pRE25_100_::*ermB*; A21 - GE1::pRE25_100_::Δ*ermB*; A22 - GE1::pRE25_300_::*ermB*; A23 - GE1::pRE25_300_::Δ*ermB*; A45 - 64/3::pRE25_100_::*ermB*; A46 - 64/3::pRE25_100_::Δ*ermB*; A47 - 64/3::pRE25_300_::*ermB*; A48 - 64/3::pRE25_300_::Δ*ermB*); F) Antibiotic resistance phenotype loss for pRUM (A25 - GE1::pRUM_300_::*ermB*; A50 - 64/3::pRUM_300_::*ermB*).

For pH311 (Inc18+RepA_N), the few *E. faecium* GE1 and *E. faecium* 64/3 isolates with vancomycin resistance phenotype reversion (1.0-3.0%), still carried the same plasmid backbone as the original plasmids and the *vanA* gene (3/5). However, the characterisation of Tn*1546* showed changes in its structure that might be related to the reversion of the vancomycin resistance phenotype (Supplementary Tables S2 and S3). For pH182 (RepA_N), phenotype reversion also occurred sporadically (1.0-4.0%) and was accompanied by the loss of Tn*1546* and often, by changes in plasmid size that did not involve the loss of key plasmid modules. Conversely, reversion to a vancomycin susceptible phenotype was frequent (up to 100%) for pBM4165 (Inc18+RepA_N) and occurred more rapidly in *E. faecium* GE1 (100% at 200 generations) than in *E. faecium* 64/3 (up to 92.0% at 300 generations). The phenotype reversion was accompanied by the loss of Tn*1546* without the loss of the plasmid or plasmid modules (Figure 3).

The behaviour of the prototype plasmids pRUM and pRE25 was similar to that of the Tn*1546*-carrying plasmids pH311 and pH182, respectively. Strains harbouring pRUM rarely lost the erythromycin resistant phenotype (<1.0%) in both *E. faecium* GE1 and *E. faecium* 64/3, and plasmid size and backbone were preserved in most cases. Conversely, pRE25 was less stable in *E. faecium* (plasmid loss of 7.0-27.0% in *E. faecium* GE1 and 0-17.0% in *E. faecium* 64/3). In some instances, the loss of erythromycin resistance was accompanied by plasmid loss, but in other cases the plasmid was present but with a different size due to the deletions that include one or more plasmid modules. Lastly, the prototype plasmid pIP501 had low stability in *E. faecium* backgrounds, being easily lost, particularly in *E. faecium* GE1 (68.0-96.0% *vs* 8.0-23.0% in *E. faecium* 64/3). The reversion to an erythromycin susceptible phenotype frequently coincided with plasmid loss (Figure 3).

### Whole genome analysis of evolved and non-evolved recipient strains

All sequenced evolved strains showed point mutations, deletions, duplications and possible recombination regions (both on the chromosome and plasmids) when compared with their corresponding parental non-evolved strains. In general, more mutations were detected in *E. faecium* 64/3 than in *E. faecium* GE1 (Supplementary Tables S2 and S3), suggesting that the frequency of mutations might correlate with plasmid adaptation.

Chromosomal changes were associated with disparate functions (DNA replication and repair, transposition and genome plasticity), environmental information processing, tissue adherence, transcription regulators, inorganic ion transport and metabolism, cellular and cell wall enzymes, and genetic information processing) (Supplementary Tables S2 and S3). Remarkably, most of the *E. faecium* GE1 showed an A160T mutation within an IS*L3*-like element, a 292bp deletion in the intergenic region encoding for 16S rRNA and tRNA, and a 211bp deletion within IS*6770*. Plasmid changes included point mutations, duplications, deletions of plasmid modules, and antibiotic resistance genes/genetic determinants that would explain the observed loss of vancomycin and/or erythromycin resistance phenotypes (see above).

### Changes in growth fitness in strains harbouring plasmids during serial passages

Most evolved strains carrying Tn*1546*-plasmids improve their growth rate when compared with the non-evolved strains (6% and 25%), often under antibiotic induction with vancomycin (Figure 4). Note that these plasmids are natural plasmids widely spread among strains of different areas.^1^

**Figure 4.**
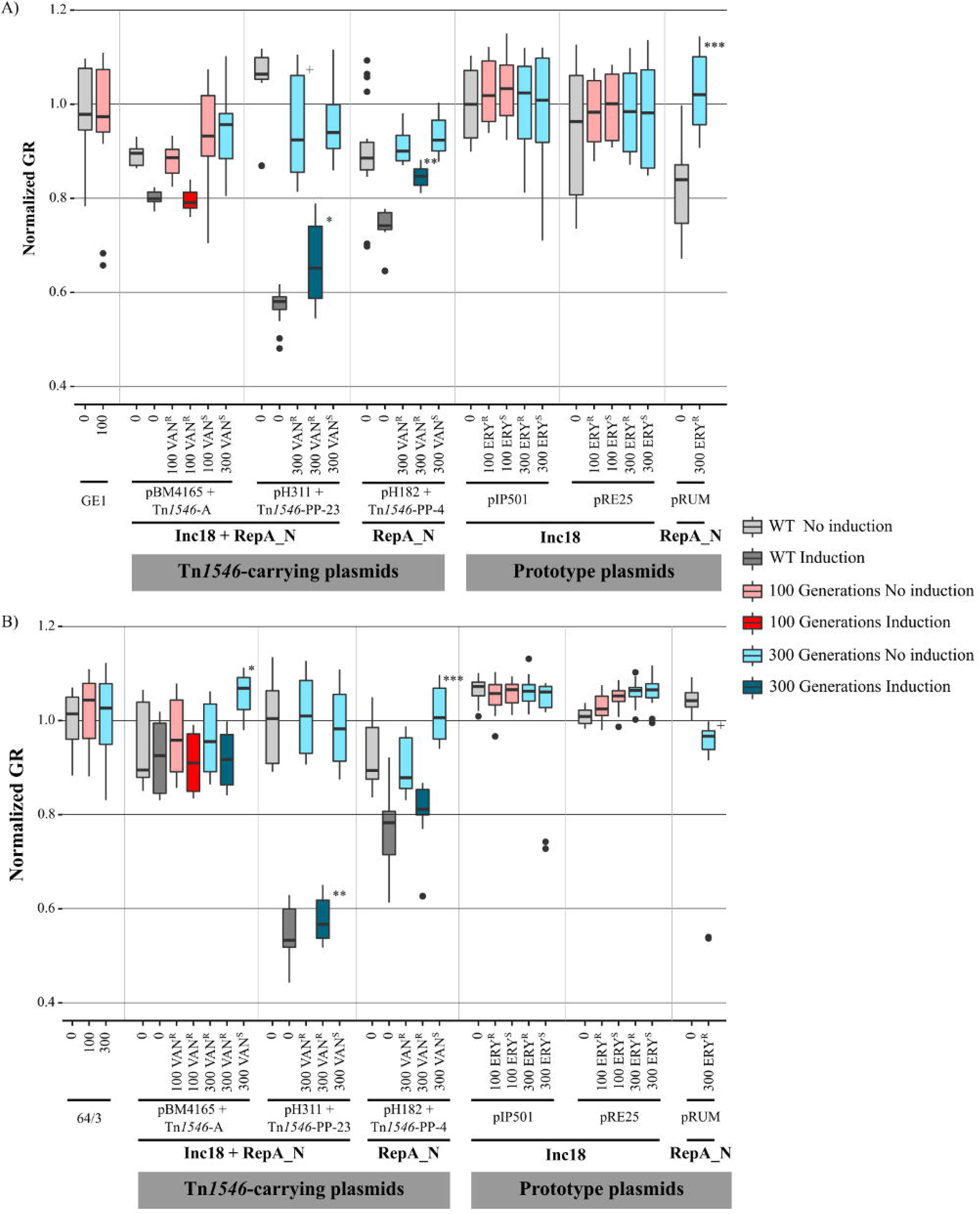
Box and whiskers plot representing the normalised growth rates of *E. faecium* Tn*1546*-*vanA*-carrying plasmids with and without vancomycin induction and prototype plasmids (pRE25, pIP501 and pRUM). A) Plasmid and Tn*1546*-*vanA* expression tested in *E. faecium* GE1. +Significantly less fit than pH311+Tn*1546*-PP-23; *Significantly more fit than induced pH311+Tn*1546*-PP-23; ** Significantly more fit than induced pH182+Tn*1546*-PP-4; *** Significantly more fit than pRUM. B) Plasmid and Tn*1546*-*vanA* expression tested in *E. faecium* 64/3. Plasmids are classified according to the replication initiation proteins plasmid family and Tn*1546* type. *Significantly more fit than pBM4165+Tn*1546*-A, pBM4165+Tn*1546*-A_100VAN^R^ and pBM4165+Tn*1546*-A_300VAN^R^; **Significantly more fit than induced pH311+Tn*1546*-PP-23; ***Significantly more fit than pH182+Tn*1546*-PP-4 and pH182+Tn*1546*-PP-4_300VAN^R^; *** Significantly less fit than pRUM. Abbreviations: WT,wild type.

### Growth fitness of representative wild-type *E. faecium* clones and clades at 37°C and 42°C

Figure S4 shows the significantly higher (p<0.01) mean growth rate of the field strains used in the study at 42°C (1.0462±0.2282) when compared with 37°C (1.024737±0.1631). A further significant difference (p<0.0001) in growth rates was observed between the isolates of the three *E. faecium* genomic clades, with Clade B having the best mean growth rate at both 37°C and 42°C. At 37°C, the mean growth rate within clades A2 and A1 varied greatly (p<0.05) (Figure S5). VSEfm strains showed a better growth rate than VREfm strains (p<0.0001), and AREfm had a significantly (p<0.0001) lower growth rate than ASEfm within VSEfm and VREfm groups (Figure S6).

## DISCUSSION

The fitness cost and stability of natural widespread plasmids carrying *vanA* (Tn*1546*) vary greatly with the clonal background of *E. faecium* and *E. faecalis* species, although, certain consistent patterns can help explain their successful distribution in humans and animals since they were first reported.

The fitness analysis of vancomycin-resistant plasmids in *E. faecium* remains mostly unexplored. A previous study suggested relative stability for these plasmids but with a tendency to plasmid loss that was not observed for most Tn*1546*-carrying plasmids tested in our study.^8^ Conversely, most Tn*1546*-*vanA*-carrying plasmids with high fitness cost were stably maintained in the bacterial population in the absence of selective pressure, eventually increasing the fitness in the presence of antibiotics. A number of these plasmids carry known, highly effective TA systems, which might account for their stability even when the vancomycin resistance genotype/phenotype was lost. However, the stability of the prototype Inc18 or pRUM plasmids carrying such TA was lower than that of the Tn*1546*-*vanA* plasmids (chimeras of Inc18 and pRUM or other RepA_N plasmids^1^), which were lost despite the low fitness cost they impose. The presence of a mixed population of carriers of Inc18 plasmid prototypes (Host::pIP501 and Host::ΔpIP501) differing in the fitness between the populations could result in cells without pIP501 that outgrow those with pIP501.^33^ Such VRE/VSE population heterogeneity was not detected in our study, but only one colony from those obtained after serial passages was the founder in the growth fitness experiments. We cannot rule out the presence of other evolved populations with increased fitness.

Most of the changes observed in the *E. faecium* evolved strains were chromosomal compensatory mutations associated with host-plasmid adaptation as previously reported^33,34^ and included DNA replication and repair mechanisms suggesting that increased mutation rates might be relevant for a rapid host-plasmid adaptation.^35,36^ Alterations in the global transcription regulation (leading to higher transcription rates in the non-evolved strains and back to normal levels in the evolved strains) have also been reported for streptomycin resistant mutants of *Salmonella* Typhimurium and in a *Pseudomonas aeruginosa* strain carrying a mercury resistance plasmid as a consequence of resistance adaptation.^36–38^ Changes found in carbohydrate and amino acid metabolism, several phosphotransferase systems and inorganic ion transport systems might be related to the metabolic burden imposed by the plasmid, leading to higher nutrient uptake and processing.^36^ It is of note that although the same functions were affected in both *E. faecium* GE1 and *E. faecium* 64/3, the mutations found in the strains of different backgrounds carrying the same plasmids were different, indicating that even when confronted with the same plasmid, the hosts might have alternative and unique compensatory evolutionary pathways.^37^

The number of Tn*1546* variants analysed here, each located in different but widespread plasmids, preclude an accurate comparison of the cost of transposon variants in different plasmid backgrounds, but also reflect the natural diversity found in hospitals.^1^ The higher fitness cost of Tn*1546* variants with indels compared with strains carrying a complete copy of Tn*1546* might explain the lack of stability of multiple Tn*1546* variants and the selection and spread of a very few variants.^39^ Although reversion of the vancomycin resistant phenotype was observed for some plasmids due to partial or complete deletions of the Tn*1546,* the stability of the plasmid backbones was documented. A result that would suggest the persistence of successful plasmid entities with ability to acquire antibiotic resistance determinants including Tn*1546* elements in independent as suggested by epidemiological studies that reflect the association of a plethora of *vanA*-Tn*1546* variants with successful plasmid chimeras.^1^ Deletion of plasmid modules and, to a lesser extent, certain mutations related with plasmid replication were observed, which have been previously documented for Inc18 and RepA_N plasmid families.^15^

The presence of a higher number of acquired antibiotic resistance and virulence genes in strains of clade A1 agrees with previous studies.^4,5,40,41^ These differences might also account, to a certain extent, for the fitness differences between clades, with more susceptible clade B having a better fitness than the multidrug-resistant clade A1.^9–11,42–44^ The differences in fitness among clades were maintained at 42°C, a temperature at which *E. faecium* appears to have significantly better fitness, probably related to the lifestyle of this species, which can colonise a host with a wide range of temperatures.^4,45^ This study found significant differences in fitness between VSE and VRE *E. faecium*. Nevertheless, the fact that VRE were found to be less fit that VSE might not be directly related to the presence of Tn*1546*, as Tn*1546* is only expressed in the presence of an inducer, or by the presence of Tn*1546*-carrying plasmids as these seem to be highly adapted to certain *E. faecium* clonal backgrounds.^10,11^ Moreover, the presence of ampicillin-resistance in VREfm and VSEfm seems to greatly reduce the fitness of these strains, indicating that selective pressure imposed by various β-lactam antibiotics extensively used in hospitals might be important for maintaining ampicillin-resistant strains, associated to specific *E. faecium* clades and justifying their association almost exclusively with the nosocomial setting.^46,47^ Moreover, other antibiotics more frequently used in hospitals, as ticarcillin or piperacillin-tazobactam, ceftriaxone and cefotetan, are excreted at high concentrations in bile, and thus reaching the upper gut improves colonisation by ampicillin-resistant clade A1 *E. faecium.*^48^

In summary, the *E. faecium* host specificity of Tn*1546*-carrying plasmids might explain the association of vancomycin resistance with this species. Although the majority of Tn*1546-*carrying plasmids studied seem to impose a significant burden on the host strain, their stability in the absence of selective pressure and their high conjugation rates between *E. faecium* strains might explain their maintenance and spread among *E. faecium* populations, as is expected given the accumulated knowledge, perpetuating the presence of VRE strains worldwide.^49^ Plasmid chimeras increase the robustness of enterococcal populations in adapting to environmental challenges including the selection pressures of antibiotics.

## TRANSPARENCY DECLARATION

None do declare.

## SUPPLEMENTARY TABLES AND FIGURES

**Table S1. Epidemiological, virulence and mobilome characteristics of wild type strains used in this study.** In plasmid content the plasmid in bold is the one carrying Tn*1546* transposon. * in bold - statistically significant; green in T=42°C GR indicates a higher fitness of the strain at 42°C compared with 37°C; Red in T=42°C GR indicates a fitness loss of the strain at 42°C compared with 37°C. Replication initiation proteins, relaxases and toxin-antitoxin systems were determined as previously reported by Freitas *et al.*^1^ Abbreviations: AbR, antibiotic resistance; VF, virulence factors; ND, Not determined.

**Table S2. Mutations in evolved strains in *E. faecium* GE1RF background**. Each colour represents a pair of evolved strains. We employed the Uniprot and KEGG databases to assess the protein functions of mutated proteins. Abbreviations: AbR, antibiotic resistance; CH, chromosome; ERY, erythromycin; P, plasmid; S, susceptible; R, resistant; VAN, vancomycin.

**Table S3. Mutations in evolved strains in *E. faecium* 64/3 background**. Each colour represents a pair of evolved strains. We employed the Uniprot and KEGG databases to assess the protein functions of mutated proteins. Abbreviations: AbR, antibiotic resistance; CH, chromosome; ERY, erythromycin; P, plasmid; S, susceptible; R, resistant; VAN, vancomycin.

**Figure S1. List of sequenced evolved and non-evolved strains with antibiotic resistance (genotypical and phenotypical) and plasmid characteristics**. Abbreviations: AbR, antibiotic resistance; CH, Chromosome; Tn, Transposon Tn*1546*

**Figure S2. PLACNET analysis of non-evolved strains in *E. faecium* GE1RF**. A) PLACNET of GE1RF; B) PLACNET analysis of GE1RF::pH311; C) PLACNET analysis of GE1RF::pH182; D) PLACNET analysis of GE1RF::pBM4165; E) PLACNET analysis of GE1RF::pIP501; F) PLACNET analysis of GE1RF::pRE25; G) PLACNET analysis of GE1RF::pRUM.

**Figure S3. PLACNET analysis of non-evolved strains in *E. faecium* 64/3**. A) PLACNET of 64/3; B) PLACNET analysis of 64/3::pH311; C) PLACNET analysis of 64/3::pBM4165; D) PLACNET analysis of 64/3::pIP501; E) PLACNET analysis of 64/3::pRUM.

**Figure S4. Box and whiskers plot of normalised growth rates at 37°C and 42°C of all *E. faecium* wild type strains employed in this study**. Abbreviations: GR, growth rate.

**Figure S5. Normalised Growth Rate analysis of *E. faecium* clades at 37°C and 42°C**. A) Box and whiskers plot of the different *E. faecium* clades (A1, A2 and B) at 37°C and 42°C. B) Summary table of Growth Rates and Genome size by clade and Bayesian Analysis of Population structure (BAPS) groups. Abbreviations: GR, growth rate; RGR, Relative Growth Rate.

**Figure S6. Normalised Growth Rates of *E. faecium* wild type strains according to vancomycin and ampicillin resistance.** A) Box and whiskers plot representing growth rates of *E. faecium* wild type strains according to vancomycin and ampicillin resistance; B) Summary table of growth rates and Genome size by vancomycin and ampicillin resistance. Abbreviations: AmpR, ampicillin resistance; AmpS, ampicillin susceptibility; GR, growth rate; RGR, relative growth rate; VanR, vancomycin resistance; VanS, vancomycin susceptibility.

