## Supplementary Figure for "Fitness cost of vancomycin-resistant *Enterococcus faecium* plasmids associated with hospital infection outbreaks"

| Name | Plasmid | Generation | Recipient Strain | PFGE | AbR phenotype | | AbR genotype | | Plasmid | RepA_N | | | Inc18 | | | | | Fitness Cost | |
|---|---|---|---|---|---|---|---|---|---|---|---|---|---|---|---|---|---|---|---|
| | | | | | | | | | | rep17 | rel3 | TA1 | rep1 | rep2 | rel6 | rel7 | TA2 | | |
| | | | | | VAN | ERY | *vanA* | *ermB* | size | pRUM | pRUM | Axe | Inc18 | pRE25 | pEF1 | pRE25 | ωεζ | PL | Tn*1546* |
| A1 | - | 0 | GE1 | AS-A | - | - | - | - | - | | | | CH | | | | | - | - |
| A2 | - | 100 | GE1 | AS-A | - | - | - | - | - | | | | CH | | | | | - | - |
| A26 | - | 0 | 64/3 | AS-C | - | - | - | - | - | | | | | | | | | - | - |
| A27 | - | 100 | 64/3 | AS-C | - | - | - | - | - | | | | | | | | | - | - |
| A28 | - | 300 | 64/3 | AS-C | - | - | - | - | - | | | | | | | | | - | - |
| A4 | pH311 | 0 | GE1 | AS-A | R | R | | | 50 | | | | | | | | | - | - |
| A5 | pH311 | 300 | GE1 | AS-A | R | R | | | 50 | | | | | | | | | 9.4% | -16.5% |
| A6 | pH311 | 300 | GE1 | AS-A | S | R | | | 50 | | | | | | | | | 9.1% | - |
| A29 | pH311 | 0 | 64/3 | AS-C | R | R | | | 50 | | | | | | | | | - | - |
| A30 | pH311 | 300 | 64/3 | AS-C | R | R | | | 50 | | | | | | | | | -1.9% | -6.1% |
| A31 | pH311 | 300 | 64/3 | AS-C | S | R | - | | 50, 200 | | | | | | | | | 0.9% | - |
| A7 | pH182 | 0 | GE1 | AS-A | R | R | | | 90 | | | | CH | | | | | - | - |
| A8 | pH182 | 300 | GE1 | AS-A | R | R | | | 90 | | | | CH | | | | | -1.0% | -14.2% |
| A9 | pH182 | 300 | GE1 | AS-A | S | S | - | - | 70 | | | | CH | | | | | -3.7% | - |
| A32 | pH182 | 0 | 64/3 | AS-C | R | R | | | 90 | | | | | | | | | - | - |
| A33 | pH182 | 300 | 64/3 | AS-C | R | R | | | 90 | | | | | | | | | 2.3% | -3.8% |
| A34 | pH182 | 300 | 64/3 | AS-C | S | S | - | - | 60 | | | | | | | | | -10.3% | - |
| A10 | pBM4165 | 0 | GE1 | AS-A | R | R | | | 40, 60 | | | | | | | | | - | - |
| A11 | pBM4165 | 100 | GE1 | AS-A | R | R | | | 40, 60 | | | | | | | | | 1.5% | 0.6% |
| A12 | pBM4165 | 100 | GE1 | AS-A | S | R | | | 30, 60 | | | | | | | | | -5.2% | - |
| A13 | pBM4165 | 300 | GE1 | AS-A | S | S | - | - | 60 | | | | | | | | | -5.6% | - |
| A35 | pBM4165 | 0 | 64/3 | AS-C | R | R | | | 40, 60 | | | | | | | | | - | - |
| A36 | pBM4165 | 100 | 64/3 | AS-C | R | R | | | 40, 60 | | | | | | | | | -1.8% | 1.4% |
| A37 | pBM4165 | 300 | 64/3 | AS-C | R | R | | | 40, 60 | | | | | | | | | 1.2% | 0.7% |
| A38 | pBM4165 | 300 | 64/3 | AS-C | S | S | - | - | 60 | | | | | | | | | -11.4% | - |
| A14 | pIP501 | 0 | GE1 | AS-A | S | R | - | | 30 | | | | | | | | | - | - |
| A15 | pIP501 | 100 | GE1 | AS-A | S | R | - | | 30 | | | | | | | | | -2.9% | - |
| A16 | pIP501 | 100 | GE1 | AS-A.2 | S | S | - | - | - | | | | CH | | | | | -3.3% | - |
| A17 | pIP501 | 300 | GE1 | AS-A | S | R | - | | 30 | | | | | | | | | 0.2% | - |
| A18 | pIP501 | 300 | GE1 | AS-A | S | S | - | - | - | | | | CH | | | | | 2.2% | - |
| A39 | pIP501 | 0 | 64/3 | AS-C | S | R | - | | 30 | | | | | | | | | - | - |
| A40 | pIP501 | 100 | 64/3 | AS-C | S | R | - | | 30 | | | | | | | | | 1.3% | - |
| A41 | pIP501 | 100 | 64/3 | AS-C | S | S | - | - | - | | | | | | | | | 0.7% | - |
| A42 | pIP501 | 300 | 64/3 | AS-C | S | R | - | | 30 | | | | | | | | | 0.5% | - |
| A43 | pIP501 | 300 | 64/3 | AS-C | S | S | - | - | - | | | | | | | | | 4.0% | - |
| A19 | pRE25 | 0 | GE1 | AS-A | S | R | - | | 50 | | | | | | | | | - | - |
| A20 | pRE25 | 100 | GE1 | AS-A | S | R | - | | 50 | | | | | | | | | -5.2% | - |
| A21 | pRE25 | 100 | GE1 | AS-A | S | S | - | - | - | | | | CH | | | | | -6.7% | - |
| A22 | pRE25 | 300 | GE1 | AS-A | S | R | - | | 50 | | | | | | | | | -5.5% | - |
| A23 | pRE25 | 300 | GE1 | AS-A | S | S | - | - | - | | | | CH | | | | | -4.8% | - |
| A44 | pRE25 | 0 | 64/3 | AS-C | S | R | - | | 50 | | | | | | | | | - | - |
| A45 | pRE25 | 100 | 64/3 | AS-C | S | R | - | | 50 | | | | | | | | | -2.1% | - |
| A46 | pRE25 | 100 | 64/3 | AS-C | S | S | - | - | 25 | | | | | | | | | -4.0% | - |
| A47 | pRE25 | 300 | 64/3 | AS-C | S | R | - | | 50 | | | | | | | | | -5.2% | - |
| A48 | pRE25 | 300 | 64/3 | AS-C | S | S | - | - | 30 | | | | | | | | | -5.5% | - |
| A24 | pRUM | 0 | GE1 | AS-A | S | R | - | | 30, 200 | | | | CH | | | | | - | - |
| A25 | pRUM | 300 | GE1 | AS-A | S | R | - | | 200 | | | | CH | | | | | -25.1% | - |
| A49 | pRUM | 0 | 64/3 | AS-C | S | R | - | | 30, 200 | | | | | | | | | - | - |
| A50 | pRUM | 300 | 64/3 | AS-C | S | R | - | | 200 | | | | | | | | | 11.7% | - |

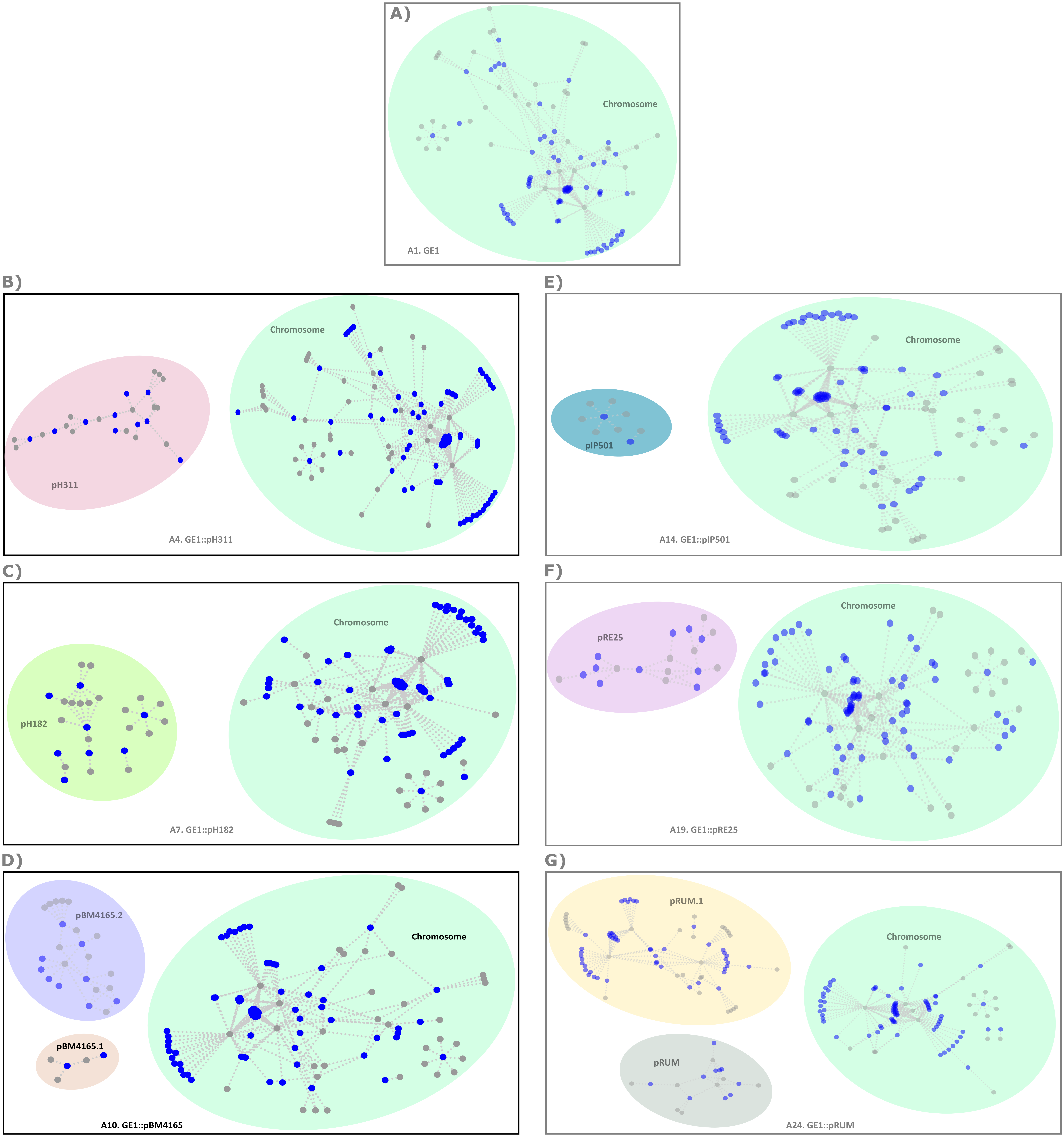
A)
Chromosome
A1. GE1
B)
pH311
Chromosome
A4. GE1::pH311
C)
pH182
Chromosome
A7. GE1::pH182
D)
pBM4165.2
pBM4165.1
Chromosome
A10. GE1::pBM4165
E)
pIP501
Chromosome
A14. GE1::pIP501
F)
pRE25
Chromosome
A19. GE1::pRE25
G)
pRUM.1
pRUM
Chromosome
A24. GE1::pRUM

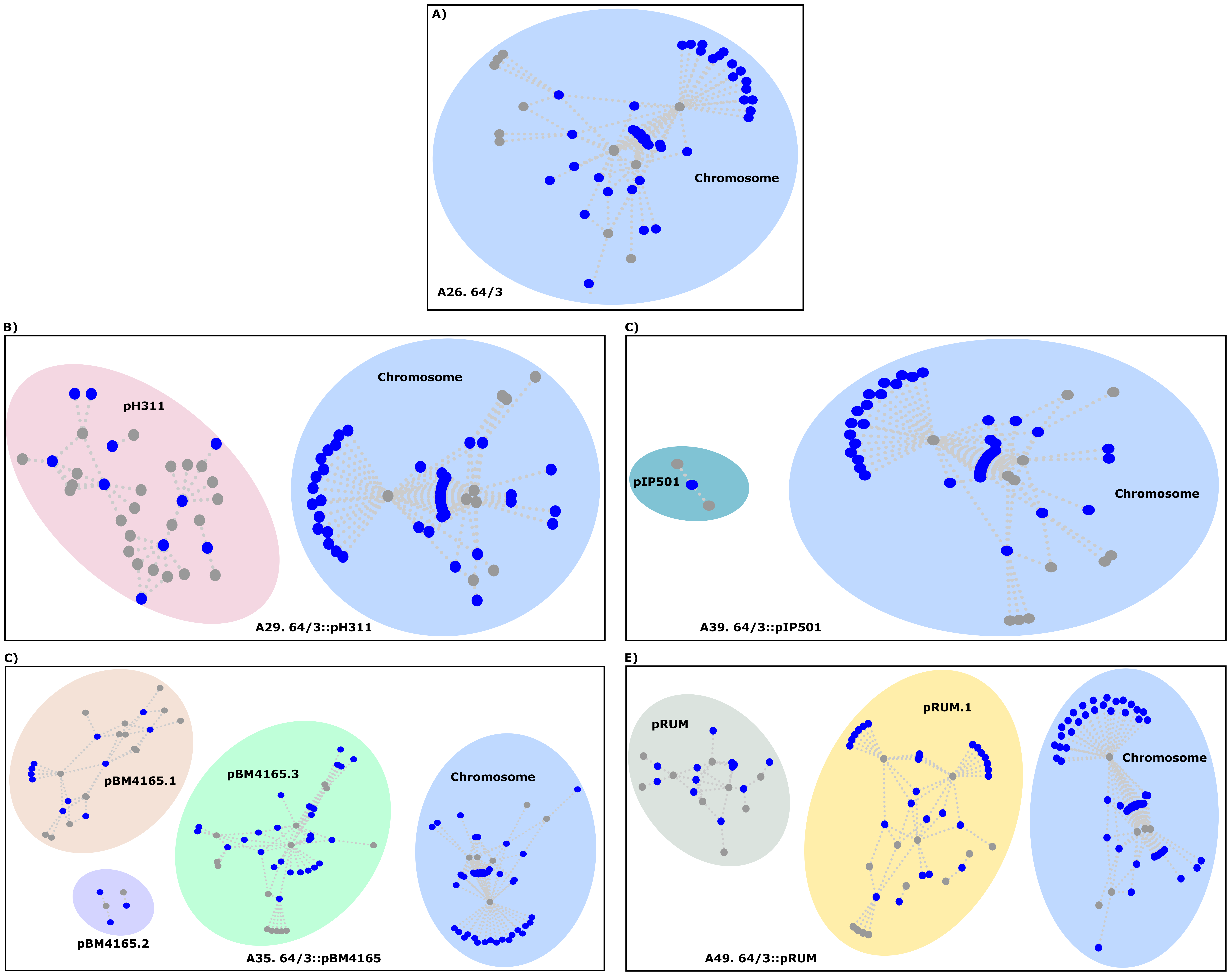

A)
Chromosome
A26. 64/3
B)
pH311
Chromosome
A29. 64/3::pH311
C)
pIP501
Chromosome
A39. 64/3::pIP501
C)
pBM4165.1
pBM4165.3
Chromosome
pBM4165.2
A35. 64/3::pBM4165
E)
pRUM
pRUM.1
Chromosome
A49. 64/3::pRUM

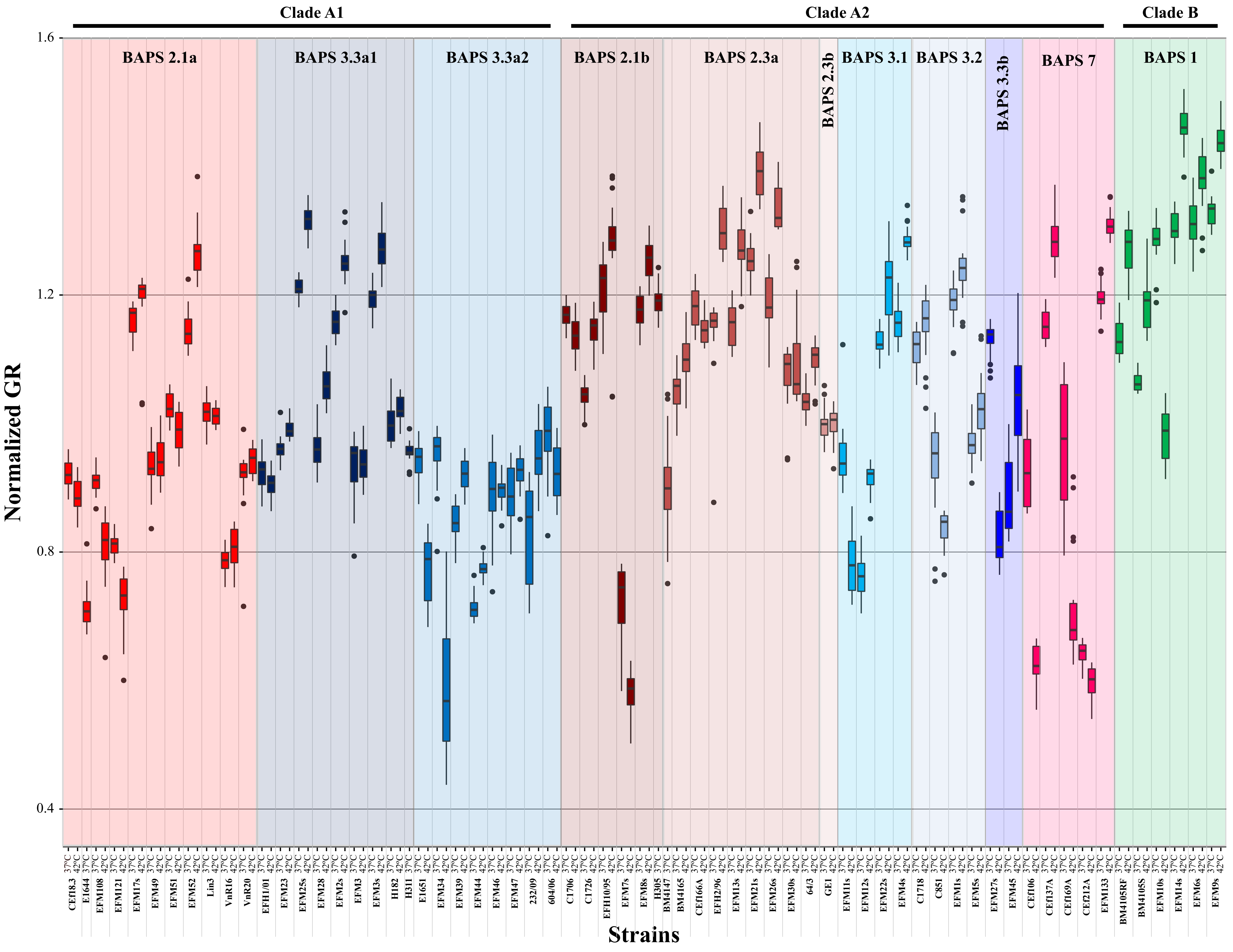

Clade A1
Clade A2
Clade B
BAPS 2.1a
BAPS 3.3a1
BAPS 3.3a2
BAPS 2.1b
BAPS 2.3a
BAPS 2.3b
BAPS 3.1
BAPS 3.2
BAPS 3.3b
BAPS 7
BAPS 1
Normalized GR
1.6
1.2
0.8
0.4
37°C
42°C
Cef18.3
E1644
EFM108
EFM121
EFM17s
EFM49
EFM51
EFM52
Lin3
VnR16
VnR20
EFH1/01
EFM23
EFM25s
EFM28
EFM2s
EFM3
EFM3s
H182
H311
E1651
EFM34
EFM39
EFM44
EFM46
EFM47
232/09
604/06
C1706
C1726
EFH10/95
EFM7s
EFM8s
H305
BM4147
BM4165
Cef166A
EFH2/96
EFM13s
EFM21s
EFM26s
EFM30s
64/3
GE1
EFM11s
EFM12s
EFM22s
EFM4s
C1718
C851
EFM1s
EFM5s
EFM27s
EFM45
Cef106
Cef137A
Cef169A
Cef212A
EFM133
BM4105RF
BM4105SS
EFM10s
EFM14s
EFM6s
EFM9s
Strains

A)

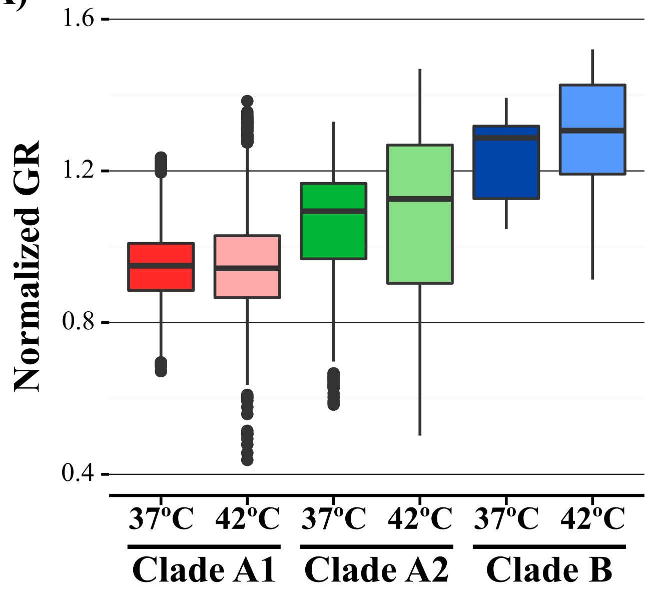


B)

| | | RGR T=37ºC | SD (±) | RGR T=42ºC | SD (±) | Genome size (Mbp) | SD (±) |
|---|---|---|---|---|---|---|---|
| **Clade B** | **BAPS 1** | 1.2378 | 0.1059 | 1.2857 | 0.1714 | 2.55 | 0.157 |
| **Clade A1** | **BAPS 2.1a** | 0.9403 | 0.1379 | 1.0334 | 0.1356 | 2.70 | 0.232 |
| | **BAPS 3.3a1** | 1.0295 | 0.1101 | 1.1085 | 0.1439 | 2.63 | 0.244 |
| | **BAPS 3.3a2** | 0.8873 | 0.0899 | 0.9408 | 0.0921 | 2.64 | 0.172 |
| | **Total** | **0.9540** | **0.1292** | **0.9672** | **0.1791** | **2.67** | **0.232** |
| **Clade A2** | **BAPS 2.1b** | 1.0835 | 0.1776 | 1.1340 | 0.2286 | 2.29 | 0.163 |
| | **BAPS 2.3a** | 1.1021 | 0.1059 | 1.1943 | 0.0962 | 2.18 | 0.089 |
| | **BAPS 3.1** | 0.9997 | 0.1626 | 1.0096 | 0.1722 | ND | ND |
| | **BAPS 3.2** | 1.0519 | 0.1136 | 1.1290 | 0.1169 | 2.41 | 0.284 |
| | **BAPS 3.3b** | 1.0014 | 0.1315 | 1.0805 | 0.2102 | 2.42 | 0.107 |
| | **BAPS 7** | 0.9758 | 0.2049 | 1.0258 | 0.2586 | 2.40 | 0.340 |
| | **Total** | **1.0506** | **0.1059** | **1.0678** | **0.2388** | **2.33** | **0.228** |

A)

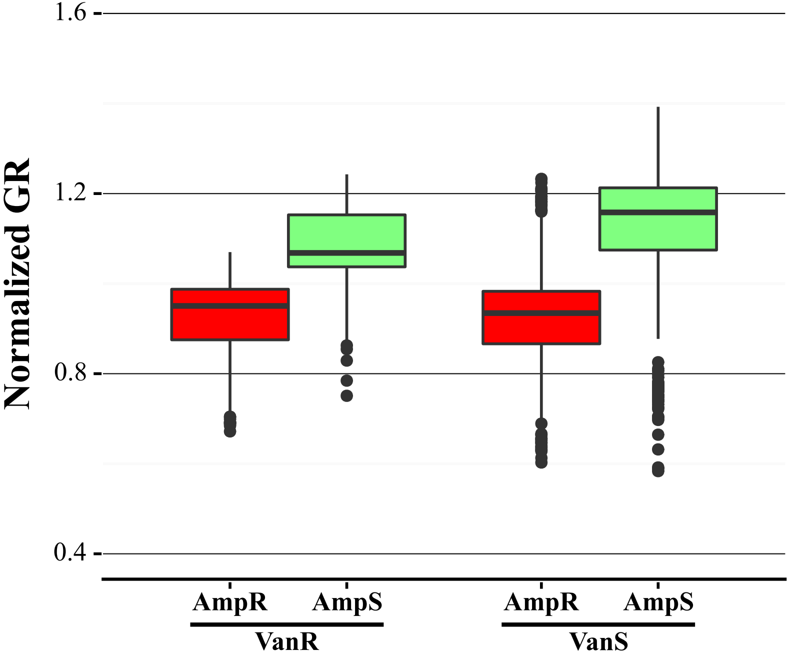


B)

| | | RGR T=37ºC | SD (±) | RGR T=42ºC | SD (±) | Genome size (Mbp) | SD (±) |
|---|---|---|---|---|---|---|---|
| **VanS** | **AmpS** | 1.1279 | 0.1477 | 1.1764 | 0.2106 | 2.40 | 0.214 |
| | **AmpR** | 0.9318 | 0.1315 | 0.0985 | 0.1959 | 2.65 | 0.222 |
| **VanR** | **AmpS** | 1.0727 | 0.0974 | 1.1337 | 0.0436 | 2.34 | 0.329 |
| | **AmpR** | 0.9182 | 0.0997 | 0.9092 | 0.0939 | 2.66 | 0.281 |
